# Alpha diversity metrics for noisy OTUs

**DOI:** 10.1101/434977

**Authors:** Robert C. Edgar, Henrik Flyvbjerg

## Abstract

Next-generation sequencing (NGS) of marker genes such as 16S ribosomal RNA is widely used to survey microbial communities. The in-sample (alpha) diversity of Operational Taxonomic Units (OTUs) is often summarized by metrics such as richness or entropy which are calculated from observed abundances, or by estimators such as Chao1 which extrapolate to unobserved OTUs. Most such measures are adopted from traditional biodiversity studies, where observational error can often be neglected. However, errors introduced by next-generation amplicon sequencing tend to induce spurious OTUs and spurious counts in OTU tables, both of which are especially prevalent at low abundances. In consequence, traditional metrics may be grossly inaccurate if they are naively applied to NGS OTU tables. In this work, we describe two novel alpha diversity estimators which are calculated from OTU abundances above a specified threshold. The singleton-free estimator (SFE) is a non-parametric estimator which is derived from a similar approach to Chao1 but extrapolates using doublet and triplet abundances rather than singletons and doublets. The octave estimator (OE) fits a log-normal distribution to non-singleton bars of an octave plot. We show that these estimators are effective under suitable conditions, but these conditions rarely apply in practice. We conclude that extrapolating to unobserved OTUs remains an open problem which is unlikely to be solved in the near future.

## Introduction

Metagenomics by next-generation sequencing (NGS) of marker genes such as 16S ribosomal RNA (rRNA) has revolutionized the study of microbial communities in environments ranging from the human body (Cho and Blaser, 2012; Pflughoeft and Versalovic, 2012) to oceans (Moran, 2015) and soils (Hartmann *et al.*, 2014). Data analysis in such studies typically assigns reads to clusters of similar sequences called Operational Taxonomic Units (OTUs). Alpha diversity, i.e. diversity of a single sample or community, is often characterized by a number (*metric*) calculated from the set of OTU abundances. For example, richness is the number of OTUs, and Shannon entropy (Shannon, 1948) is a function of the OTU frequencies. In both traditional and metagenomic studies, some groups (species or OTUs) may be missing because of insufficient observations; we will refer to this as *incomplete enumeration* to avoid over-use of the terms sample and sampling. Some metrics, notably Chao1 (Chao, 1984; Chiu *et al.*, 2014), attempt to account for unobserved OTUs by extrapolating from the observed abundances; we shall refer to such metrics as *estimators*.

### Amplicon sequencing errors and biases degrade diversity measurements

While experimental error can largely be neglected in traditional studies, metagenomic OTUs are often spurious due to sequence errors (Edgar, 2013; Huse *et al.*, 2010), which can lead to grossly incorrect estimates of diversity (Edgar, 2017a). Biases favoring or disfavoring observations of some groups are recognized in traditional biodiversity studies (for example, larger species are easier to see), but are often considered to be inconsequential in communities of similar organisms (e.g., birds). In marker gene metagenomics, biases due to mismatches with PCR primers and variations in gene copy number are more severe, causing abundances of gene sequences in the reads to have low correlation with species abundances (Edgar, 2017b).

### Abundance filtering

Incorrect sequences reported by next-generation amplicon sequencing are strongly biased to occur with low abundances, with singleton sequences and singleton OTUs in particular highly enriched for errors (Edgar, 2013). Quality and chimera filtering can reduce but not eliminate incorrect sequences (Edgar and Flyvbjerg, 2015; Edgar, 2016). Counter-intuitively, even if the rate of incorrect bases is very low after quality filtering, a large majority of unique read sequences is nevertheless likely to be incorrect, most of which will have low abundances (https://www.drive5.com/usearch/manual/tolstoy.html). While many of these bad sequences are readily recognized and can be corrected (Callahan *et al.*, 2016; Edgar, 2017c), a substantial subset is likely to induce spurious OTUs if retained (Edgar and Flyvbjerg, 2015; Edgar, 2013). Methods which aim to minimize spurious OTUs therefore discard low-abundance sequences. For example, by default UPARSE (Edgar, 2013) discards singleton reads (sequences found only once in the dataset), DADA2 (Callahan *et al.*, 2016) sets a minimum abundance of four per sample, and UNOISE2 (Edgar, 2017c) sets a minimum of eight per dataset. While these defaults have been shown to give good results on mock community tests, they may be sub-optimal for some datasets; for example, we have suggested (Edgar and Flyvbjerg, 2018) that a minimum abundance of ~100 reads per OTU per sample should be applied in the prostate cancer microbiome study of (Yow *et al.*, 2017).

### Cross-talk errors

Multiple samples are routinely multiplexed into a single run by embedding index sequences into PCR primers (Walters *et al.*, 2011; Kozich *et al.*, 2013). A *cross-talk* error occurs when a read is assigned to an incorrect sample, e.g. because of a base call error in the index sequence. Cross-talk rates of ~1% have been observed with both pyrosequencing (Carlsen *et al.*, 2012) and Illumina (Kircher *et al.*, 2012; Nelson *et al.*, 2014; Edgar, 2018). The cross-talk rate can be measured by including control samples such as mock communities or distilled water, but cross-talk errors cannot be reliably filtered from *in vivo* samples even if the rate is known (Edgar, 2018).

Most cross-talk errors probably cause minor fluctuations in observed counts at comparable levels to those caused by enumeration, and these will have little impact on diversity estimates. However, if an OTU (*x*) is in fact absent from a sample (*A*), cross-talk can cause *x* to have a spurious non-zero count for *A* in the OTU table. Spurious non-zero counts due to cross-talk can substantially inflate diversity metrics such as richness and Chao1 that are sensitive to low-abundance counts.

Consider a sample *S* and the “meta-sample” *M* composed of all samples other than *S*. *M* is likely to have higher alpha diversity than *S* and a substantially different composition. Cross-talk from *M* into *S* will then have a strong tendency to induce non-zero counts for OTUs that are in fact absent from *S* or unobserved in *S*. While the latter case is naively benign, the observed number of low-abundance OTUs in *S* is over-estimated, which may degrade extrapolation of the distribution and comparisons between samples. Thus, cross-talk will tend to systematically inflate alpha diversity and deflate beta diversity for all samples sequenced in the same run unless two conditions are satisfied that rarely apply in practice: (1) all samples have similar composition (low beta diversity) and (2) all samples are mostly or fully enumerated (have few or no unobserved OTUs). Spurious counts due to cross-talk are often greater than one (Edgar, 2018), which implies that discarding singletons per sample could mitigate cross-talk errors to some extent but may leave many spurious counts in the OTU table.

### Truncated estimators

As the above discussion shows, even the best current analysis pipelines may miss low-abundance OTUs that are present in the reads because low-abundance sequences are discarded before clustering. Also, they may produce spurious OTUs due to sequence error and cross-talk. Therefore, alpha diversity metrics that are sensitive to low-abundance counts, such as richness and Chao1, may be inaccurate and misleading. One approach to addressing this problem is to design *truncated estimators* that consider only OTU abundances above a threshold, using them to extrapolate to the full diversity of the sample. Extrapolation may be limited to inferring the number of singletons (plus any other discarded values), or it may include unobserved OTUs. To the best of our knowledge, the only previously published truncated estimator is *breakaway_nof1* (Willis, 2016), which estimates the richness of a sample without using singleton counts. The *breakaway_nof1* estimates are obtained by a least-squares fit of the parameters of a statistical model that generates frequency ratios *f*_i+1_/*f*_*i*_, where *f*_*i*_=*n*_*i*_/*N*_*obs*_, *n*_*i*_ is the observed number of OTUs with abundance *i*, and *N*_*obs*_=Σ_*i*_ *n*_*i*_ is the total number of observed OTUs.

Here, we describe two novel truncated estimators: the singleton-free estimator (SFE) and and the octave estimator (OE). SFE is a non-parametric estimator that is derived by logical extension of the derivation of Chao1. It extrapolates using doublet and triplet abundances rather than singletons and doublets. OE fits a log-normal distribution to OTU abundances above a threshold, which is set to one by default.

## Methods

### Chao1 estimator

The Chao1 estimator is calculated from the observed number of OTUs (*Nobs*), number of singletons (*n*_1_) and number of doublets (*n*_2_) as Chao1 = *N*_*obs*_ + *n*_1_^2^/2*n*_2_.

### Singleton-free estimator (SFE)

Let *n*\*_*i*_ be the estimated number of OTUs with abundance *i*, with *n*\*_0_ representing the estimated number of unobserved OTUs. Under similar assumptions to those used to derive the Chao1 formula, we obtain the following estimates (Appendix eqs. A29 and A30):

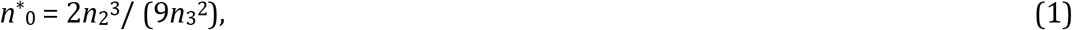

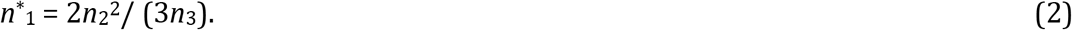

The singleton-free estimator is calculated as the observed number of non-singleton OTUs (*N*_*obs*_− *n*_1_) plus the estimated correct number of singletons and unobserved OTUs,

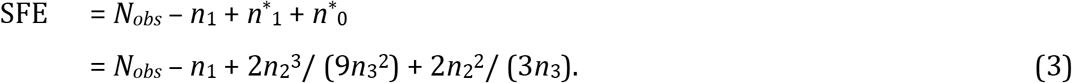

### Octave estimator (OE)

The octave estimator fits a normal distribution to an abundance histogram plotted on a log2 scale, motivated by the expectation that many abundance distributions (ADs) encountered in practice will be approximately log-normal. The log-normal model is attractive both empirically and theoretically because most observed macro-ecology ADs are well-approximated by a log-normal distribution, and by the central limit theorem a community’s AD will be log-normal if its abundances are determined by many independent random factors (May, 1975). Let *h*_*k*_ be the number of OTUs with an abundance (*r*) in the range *r*=2^*k*^, 2^*k*^+1 … 2^*k*+1^−1, using the bin boundaries of (Edgar and Flyvbjerg, 2018). We shall refer to *h*_*k*_ as the size of the *k*th bar in the histogram. A least-squares fit of a Gaussian *g*(*k*) is made to the *h*_*k*_’s, where the first *m* bins are not used in fitting with *m*=1 by default. OE is the number of OTUs, including unobserved OTUs, predicted by the fitted Gaussian, obtained as the area under the curve. The estimate with singletons included, i.e. with *m*=0, is denoted OE(0).

### Octave ratio metrics Oct1 and Oct2

In (Edgar and Flyvbjerg, 2018), we argued that an abundance distribution can be robustly extrapolated to estimate the number of unseen OTUs if, and only if, a tail towards low-abundance OTUs is visible in an octave plot. The hockey-stick (*J*-shaped) distributions that are typically found in practice cannot be reliably extrapolated towards low-abundance OTUs because no tail is visible, and the functional form of the distribution is unclear (log-normal and log-series are likely to fit well). Also, the observed *J* shape may be an artifact of spurious low-abundance counts overlaid onto the true distribution.

If, in contrast to the *J* shape, the counts in the first few bins *increase* with increasing abundance, this suggests that the tail towards low abundance is visible in the distribution. Then spurious counts are likely to be less frequent than valid counts (because otherwise, the singleton count would be high). The following metrics indicate whether sizes of low-abundance histogram bars increase or decrease as functions of abundance:

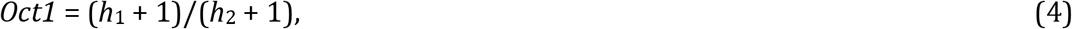

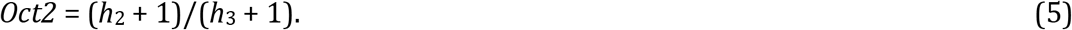

Here *h*_*i*_ is the total abundance of OTUs in bin *i*, i.e. the size of the *i*th histogram bar. We add one to the bar sizes in these definitions to avoid division by zero. If *Oct1* < 1, then the counts increase from the first to the second bar, which suggests that the tail towards low abundance may be visible in the distribution. Similarly, if *Oct2* < 1, then the same tail may be present if the sizes of bars two and three are believed to be approximately correct. These inferences follow if it is assumed that the distribution has no more than one peak and that the relevant bins are not empty; if the distribution has an anomalous shape or has been fully enumerated such that the first bins are empty then the values may be misleading. As with other inferences from summary metrics, we recommend reviewing an octave plot to aid interpretation.

### Validation on mock samples

While mock samples are essential for validating methods which generate OTUs and OTU tables, they are not suitable for validating alpha diversity metrics *per se*. If the number of observed OTUs (measured richness) differs from the known number of species, this is due to errors introduced by PCR, sequencing and OTU clustering. Mock abundances are unrealistic because diversity is very low, most or all valid biological OTUs are present in the reads, and the distributions are rarely, if ever, designed to resemble interacting ecosystems. Mock samples are therefore not suitable for testing extrapolation to unseen OTUs, and we do not consider this approach further here.

### Validation on in-vivo samples

Samples obtained *in vivo* are realistic by definition, but their diversity is not known independently of NGS. This rules out validation of extrapolation methods by comparison with a superior standard of truth. As an alternative, predictions from different estimators can be compared to each other. When estimators with distinctly different designs estimate in approximate agreement, this indicates that their estimates may be accurate. By the same reasoning, if an estimator disagrees with a consensus of other estimators, it may be less accurate.

We chose to implement this strategy using data of (Flores *et al.*, 2014), who sequenced samples obtained weekly from four human body sites (forehead, palm, tongue and gut) of 85 adults over a period of three months. This study was chosen because there are multiple body sites, enabling pair-wise comparisons between sites that were sequenced using the same protocol, and because there are many samples (3,593) which were deeply sequenced (median 42,166 reads per sample after quality filtering and discarding singletons per dataset). We generated an OTU table using the current recommended UPARSE (Edgar, 2013) protocol (https://drive5.com/usearch/manual/uparse_pipeline.html, accessed 1st August 2018). A minimum abundance of eight per dataset was imposed on the read sequences, and the OTU table was rarefied to 5,000 reads per sample to obtain a more typical read depth and reduce the number of spurious low-abundance counts due to sequence errors and cross-talk. For each metric and each pair of body sites, diversity was compared using a two-tailed Wilcoxon signed-rank test.

### Validation on simulated data

We simulated a log-normal abundance distribution by random sampling from a Gaussian probability density function with parameter values mean *μ* = 4.6 and standard deviation *σ=* 1.5, as described in (Edgar and Flyvbjerg, 2018). The size of the population was fixed at 2^15^=32,768 individuals, which generated 758 OTUs. Observed distributions were obtained by simulating 2^*k*^ reads with *k*=15, 14 … 9.

## Results

### Validation on in-vivo samples

Results in the *in vivo* samples are shown in Table 1. All metrics except Shannon entropy agree that forehead > gut > tongue, where *x* > *y* means that site *x* has greater diversity than site *y* with *P*<0.05. By entropy, forehead < gut, gut > tongue and forehead > tongue. Forehead and palm are not distinguished except by OE, which gives weak significance (*P=*0.04) to forehead > palm. On all other site pairs, all metrics agree that there is a statistically significant difference.

**Table 1.** *P*-values resulting from pair-wise site comparison. Sites are: *Fh* forehead, *Plm* palm, *Gut* gut, *Tng* tongue. Metrics are *Richness, Chao1, Shannon, SFE, OE* and *OE*(*0*) as described in the main text. Table entries are P-values, with background colors indicating the sign of the difference and its statistical significance. Color codes: y is the name of the row, x is the name of the column.

| Richness | Tng | Gut | Plm |
| --- | --- | --- | --- |
| Fh | 0 | 0 | 0.057 |
| Plm | 0 | 0 |  |
| Gut | 0 |  |  |

| Chao1 | Tng | Gut | Plm |
| --- | --- | --- | --- |
| Fh | 0 | 0 | 0.52 |
| Plm | 0 | 0 |  |
| Gut | 0 |  |  |

| Shannon | Tng | Gut | Plm |
| --- | --- | --- | --- |
| Fh | 1.3e-4 | 0 | 0.70 |
| Plm | 8.0e-5 | 0 |  |
| Gut | 0 |  |  |

| FE | Tng | Gut | Plm |
| --- | --- | --- | --- |
| Fh | 0 | 0 | 0.06 |
| Plm | 0 | 0 |  |
| Gut | 0 |  |  |

| OE | Tng | Gut | Plm |
| --- | --- | --- | --- |
| Fh | 0 | 1.8e-4 | 0.04 |
| Plm | 0 | 0 |  |
| Gut | 0 |  |  |

| OE(0) | Tng | Gut | Plm |
| --- | --- | --- | --- |
| Fh | 0 | 3.2e03 | 0.12 |
| Plm | 0 | 2.0e-5 |  |
| Gut | 0 |  |  |

$y > x, P < 0.05$ $y < x, P < 0.05$ $y < x, P \geq 0.05$ other

### Validation on simulated log-normal

Octave plots for the simulated log-normal distribution are shown in Fig. 1. We have previously proposed (Edgar and Flyvbjerg, 2018) that octave plots can be classified as complete (*C*), truncated (*T*), *J*-shaped (*J*) and anomalous, i.e. not consistent with a log-normal distribution, (*A*). With 32k and 16k reads, an approximately complete bell-curve is seen, and we would therefore classify the plots as shape *C*. With 8k to 2k reads, the curves are visibly incomplete and have shape *T*, with *J* shapes for 1k and 512 reads. If the singleton bin is disregarded, then no peak is visible with 2k or 4k reads, and these would then be classified as *J*. Metrics are reported in Table 2. These confirm the results we would intuitively expect: when a peak is visible, more than half of the diversity has been enumerated and extrapolation is reasonable. When no peak is visible in the considered bins, then extrapolation is not well-supported by the data and the predictions are less accurate. For these shapes, the *Oct1* and *Oct2* metrics indicate whether the tail is visible when singletons are included or excluded, respectively.

**Table 2.** Metric values for the simulated log-normal distribution. The correct total number of OTUs, including any unobserved, is 758. Richness is the number of OTUs which appear in the simulated reads. Metric values within 10% of the true population richness (758) are indicated by dark blue shading, within 25% by light blue shading, otherwise by light gray shading. Values of Oct1 and Oct2 are shaded light green if < 1, light orange if > 1. Notice that SFE has accuracy comparable to Chao1 when Oct2 < 1, and that both SFE and Chao1 are unreliable (further than 25% from the correct value) when Oct2 < 1.

| *Reads* | *Richness* | *Chao1* | *SFE* | *OE* | *OE(0)* | *Oct1* | *Oct2* |
| --- | --- | --- | --- | --- | --- | --- | --- |
| 32,768 | **758** | **759.0** | **758.8** | 893.0 | 892.0 | 0.250 | 0.267 |
| 16,384 | **747** | **755.2** | **740.9** | 626.0 | 933.0 | 0.324 | 0.496 |
| 8,192 | **719** | **738.4** | **723.4** | 888.0 | 897.0 | 0.374 | 0.671 |
| 4,096 | 657 | **716.0** | **721.3** | 586.0 | 586.0 | 0.615 | 0.965 |
| 2,048 | 559 | 678.4 | 592.8 | 660.0 | 545.0 | 0.872 | 1.47 |
| 1,024 | 428 | 646.6 | 474.5 | 467.0 | 467.0 | 1.45 | 2.38 |
| 512 | 297 | 558.3 | 254.8 | 315.0 | 262.0 | 1.93 | 3.76 |

**Figure 1.**
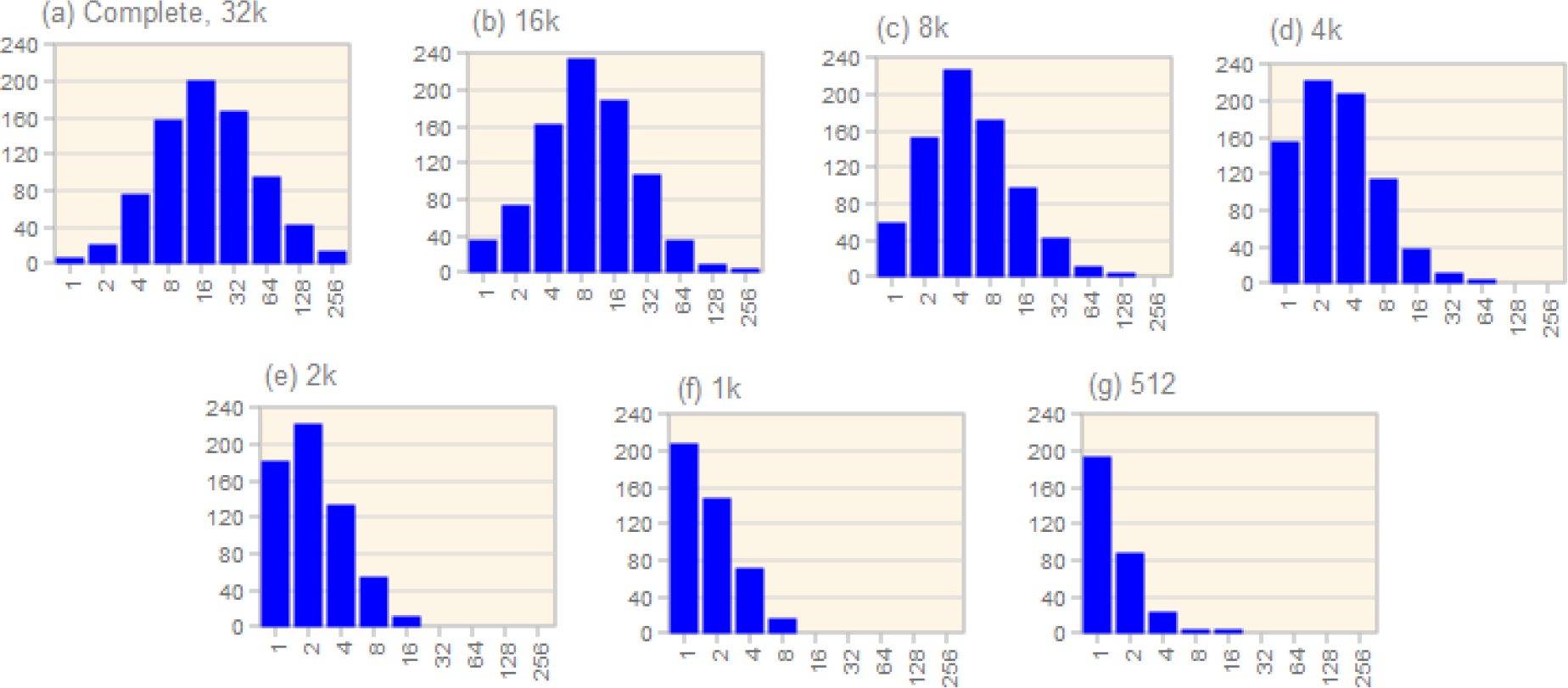
Octave plots of the simulated log-normal distribution. Panel (a) shows the complete distribution with 32k reads. The remaining panels show simulated observed distributions with from 16k reads (b) to 512 reads (g).

## Discussion

Here, we have proposed and implemented two approaches for validating alpha diversity estimators: comparing metrics on *in vivo* data, and *in-silico* simulation. Neither of these approaches is adequate to show that accurate results will be achieved in practice. On *in vivo* data, different metrics could agree due to systematic errors in the data. Simulation requires realistic models of the true diversity of the community and of errors in the data, but neither the true diversity of representative microbial communities nor amplicon sequencing errors are currently understood well enough to support the design of a convincing simulation.

Instead, we chose to implement an explicitly idealized simulation where the true distribution is log-normal and there are no simulated errors. Our results show that even in this idealized scenario, it is not possible to reliably extrapolate to the full diversity of a sample when the observed distribution is *J*-shaped, as commonly found in practice. This issue therefore applies when observational error can reasonably be neglected, as often applies in traditional biodiversity studies of macro-organisms. With microbial metagenomics by next-generation amplicon sequencing, the problem of extrapolating from a severely truncated observed distribution is exacerbated by experimental errors which are increasingly prevalent at low abundances. As a result, depending on which error suppression strategies are employed by the data analysis, an OTU table may strongly over-or under-represent low-abundance counts, and these biases cannot be satisfactorily corrected. Since accurate low-abundance counts, especially singletons, are essential for accurate extrapolation of a substantially truncated distribution, we believe that alpha diversity estimators which attempt to extrapolate are inappropriate for NGS microbial OTUs, especially Chao1. Truncated estimators such as *breakaway_nof1*, *SFE* and *OE* could be appropriate under suitable conditions, i.e. (a) abundances > 1 are known to be accurate, and also (b) the tail of the distribution is visible. However, these conditions rarely apply in practice, and we therefore believe that it is generally not possible to make reliable estimates of the full diversity of a sample from observed microbial OTUs.

## APPENDIX

## A Derivation of the singleton-free estimator

Thirty years after Chao derived the estimator of species richness that became known as Chao1 (Chao, 1984), she and coauthors derived the same estimator again in a new, easier, and much more elegant manner (Chui *et al.*, 2014). Here we generalize the latter derivation to derive an estimator of species richness that does not use the observed number of singletons, which are species (or OTUs) that are observed only once in the sample that estimation is based upon.

## A.1 Notation, definitions, and their simple relations

Table 1 summarizes the following notation: We consider a population consisting of an unknown number,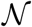, of individuals. This population is composed of an unknown number, 𝒮, of distinct species. We refer to these distinct species by distinct labels, which we choose to be the integers from 1 to *S*. Thus, when we refer to the s’th species, we mean species number *s*, and no rank is associated with *s*. It is just a label.

Now consider an attempt to characterize and enumerate the number of individuals and species in a sample from the population. Let *k*_*s*_ denote the number of individuals from the s’th species or OTU found in the sample. Let *N* denote the total number of individuals found in this enumeration. Obviously,

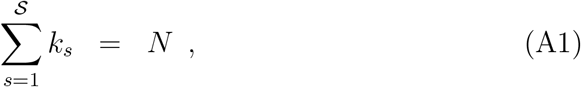

where, for some species *s*, possibly *k*_*s*_ = 0.

For notational convenience, we use Kronecker’s delta-function on the integers and the so-called *ℓ*_1_-norm for vectors,

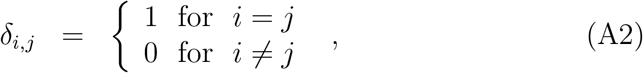

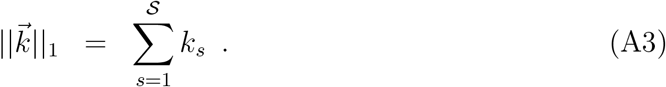

**Table A1.**
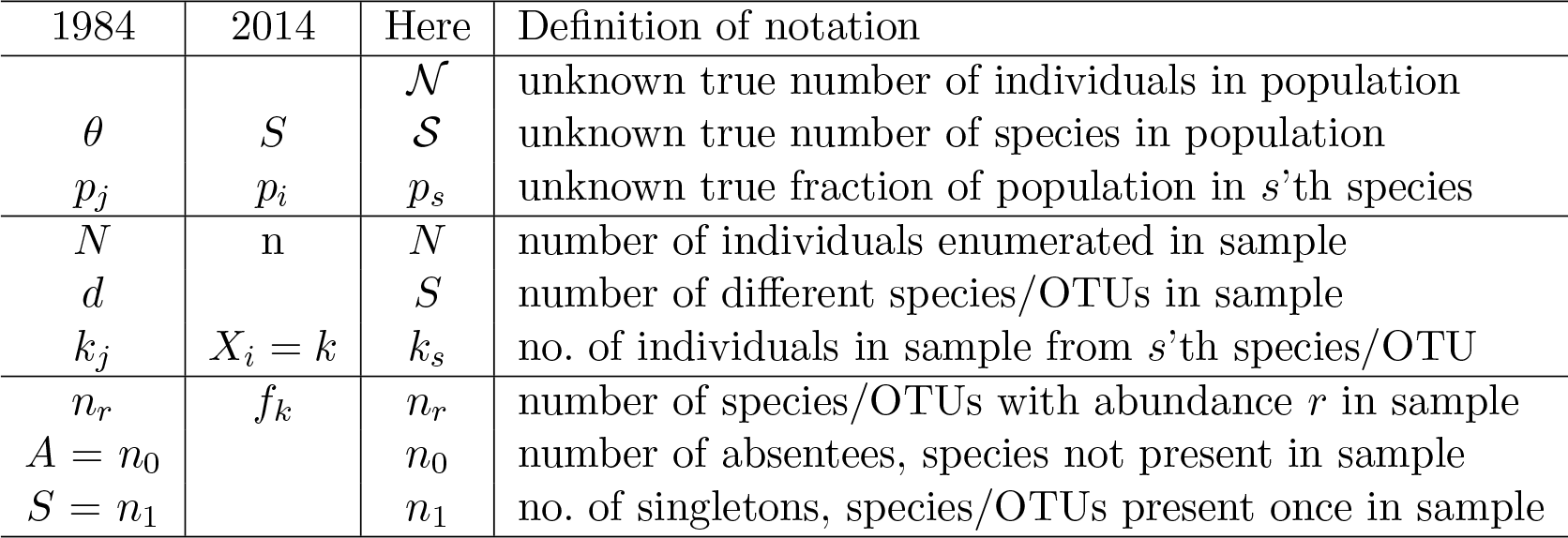
Notation used. First column: used in (Chao, 1984). Second column: used in (Chui et al., 2014). Third columns: used here.

With that, we can write any sum over all values of 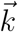 that satisfy ‖*k*‖_1_ = *N* simply as the sum over all values of 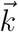 by including a factor, Kroneker’s delta, that is zero except where ‖*k*‖_1_ = *N*, where that factor equals one,

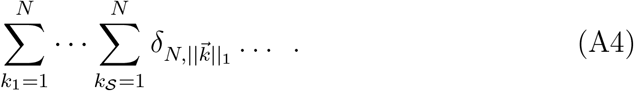

Another notational convenience is a shorthand notation for the multinomial coefficient,

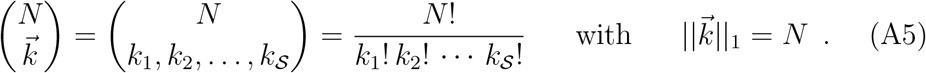

Let *n*_*r*_ denote the number of elements in 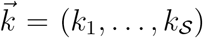 that have the value *r*. Phrased differently: *n*_*r*_ denotes the number of different species that occur with abundance *r* in the sample. Obviously, *r* is an integer with possible values in {0,1,…,*N*}, and

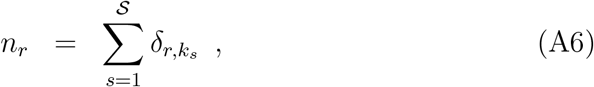

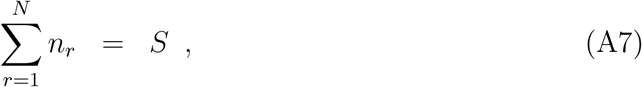

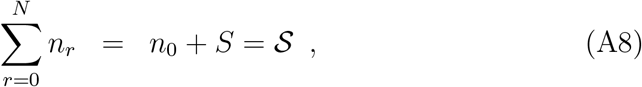

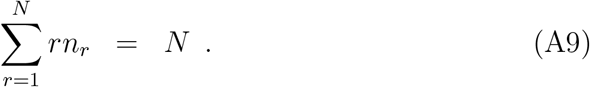

Let *p*_*s*_ denote the fraction of individuals in the population that belong to the *s*’th species. Then *p*_*s*_ is also the probability that an individual selected at random from the population belongs to the *s*’th species when any individual is equally likely to be selected. The vector 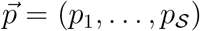 is unknown, but obviously

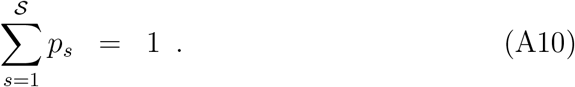

## A.2 Probabilities and expected values

## A.2.1 The numbers 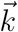 of individuals representing each species in a sample were drawn from a multinomial distribution

From Eq. (A10) follows that the number one can be written as a homogenous polynomial of degree *N* in the elements of 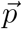,

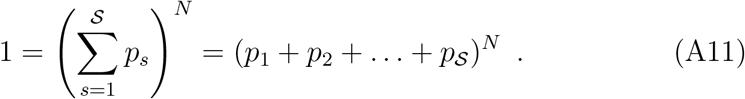

Written out without using the commutative law, each of the *S*×*N* terms that result represents the result of a sampling one individual from the population and the probability of the sample that resulted. The last statement is correct only for sampling with replacement, but is also an excellent approximation for the typical case of 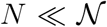, since in that case 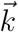 resulting from samling without replacement differs negligibly from 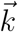 resulting from sampling with replacement.

We wish to estimate the value of, and to that end we are not interested in the order in which the individual member of the sample entered the sample in the sampling process. As a step towards estimating *S*, we are only interested in the number *k*_*s*_ of individuals in the sample belonging to species s, for *S* all species. The vector 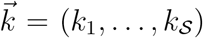 satisfying Eq. (A1) describes this composition of the sample, and the probability for its occurrence is the corresponding term in the result of applying the commutative law to the polynomial above,

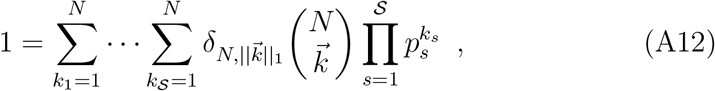

which term is the multinomial probability

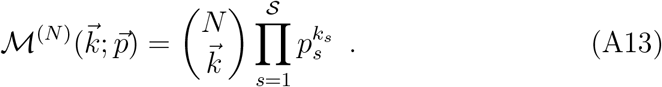

This probability distribution is obviously normalized to one.

## A.2.2 Expected value *E*[*n*_*r*_] of number of species, *n*_*r*_, of abundance *r* in sample

From Eq. (A6) follows that

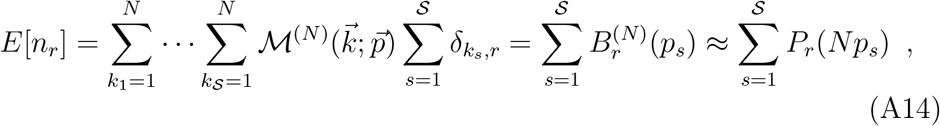

where

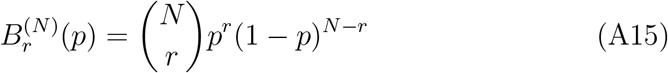

is the binomial distribution, and

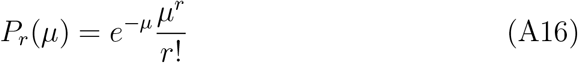

is the Poisson distribution. When *p*_*s*_ ≪1, the Poisson distribution is an excellent approximation to the binomial distribution.

## A.3 Derivation of singelton-free estimator

Define the 𝒮-dimensional vector 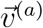 as follows,

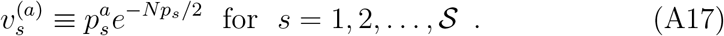

Then, using Eqs. (A14) and (A16),

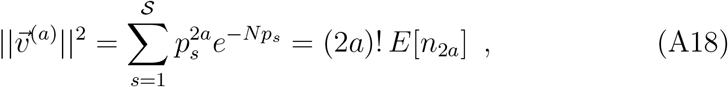

and

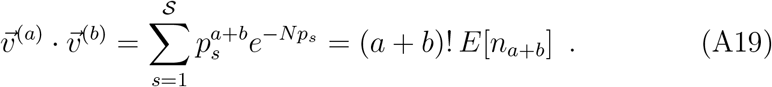

Cauchy-Schwartz’s inequality,

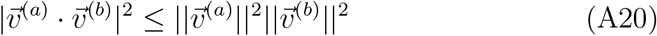

then reads

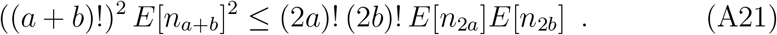

Note that if all *p*_*s*_ are equal, *p*_*s*_ = *S*^−1^, and the two sides in Cauchy-Schwartz’s inequality are equal. In this case the inequality in Eq. (A21) reduces to equality, and the approximation becomes exact. For this reason, the estimates derived in (Chui et al., 2014) are called “almost exact” there.

## A.3.1 Chao1

For *a* = 0 and *b* = 1,

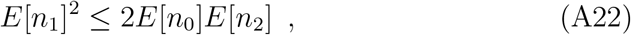

and hence

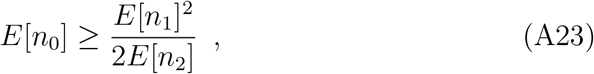

which for large values of *n*_1_ and *n*_2_ is well approximated by replacing expected values with observed values,

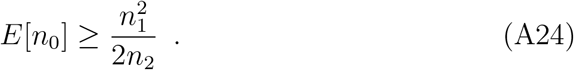

Note that the inequality in this last expression may not hold. We cannot be sure it does for any given case of observed values for *n*_1_ and *n*_2_. However, if this estimate gives a large value for *E*[*n*_0_], we can further approximate *E*[*n*_0_] with *n*_0_,

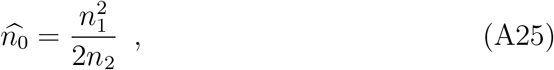

and then have

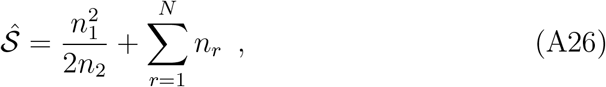

which is Chao1.

## A.3.2 Singleton-free estimator

For *a* = 1/2 and *b* = 3/2,

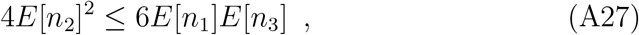

and hence

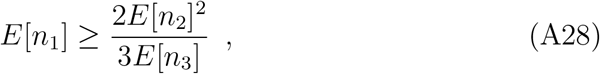

which for large values of *n*_2_, and *n*_3_ is well approximated by replacing expected values with observed values,

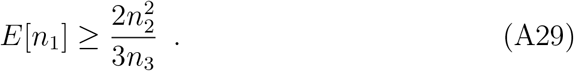

Combining Eqs. (A23) and (A28), we have

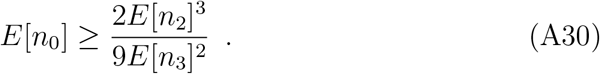

This being the result of combining two inequalities, we expect the estimator

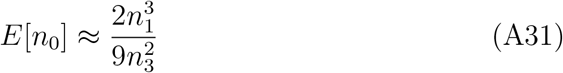

to be less accurate than Eq. (A24).

We conclude that 𝒮 is estimated by

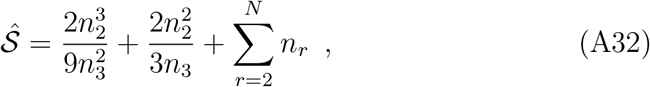

and since *n*_1_ does not occur in it, we call it singleton-free.

